# The START domain potentiates HD-ZIPIII transcriptional activity

**DOI:** 10.1101/2022.10.08.511209

**Authors:** Aman Y. Husbands, Antje Feller, Vasudha Aggarwal, Courtney E. Dresden, Ashton S. Holub, Taekjip Ha, Marja C.P. Timmermans

## Abstract

HD-ZIPIII transcription factors (TFs) were repeatedly deployed over 725 million years of evolution to regulate central developmental innovations. The START domain of this pivotal class of developmental regulators was recognized over twenty years ago, but its putative ligands and functional contributions remain unknown. Here, we demonstrate that the START domain promotes HD-ZIPIII TF homodimerization and increases transcriptional potency. Effects on transcriptional output can be ported onto heterologous TFs, consistent with principles of evolution via domain capture. We also show the START domain binds several species of phospholipids, and that mutations in conserved residues predicted to affect either ligand binding, or its downstream readout, abolish HD-ZIPIII DNA-binding competence. Our data present a model in which the START domain potentiates transcriptional activity and uses ligand-induced conformational change to render HD-ZIPIII dimers competent to bind DNA. These findings resolve a long-standing mystery in plant development and highlight the flexible and diverse regulatory potential coded within this widely distributed evolutionary module.

## Introduction

Development of multicellular organisms requires the precise control of transcription factor (TF) inputs into their gene regulatory networks. As such, the activity of TFs is highly regulated, often integrating distinct mechanisms across multiple regulatory levels to impact developmental outcomes. In plants, this is exemplified by CLASS III HOMEODOMAIN-LEUCINE ZIPPER (HD-ZIPIII) proteins, an ancient TF family that arose after the divergence of unicellular Chlorophyta but before the emergence of Streptophyte algae and land plants over 725 million years ago^*1–3*^. HD-ZIPIII TFs have been repeatedly coopted throughout plant evolution to regulate key developmental advances ^*4–13*^. For instance, in *Arabidopsis thaliana*, HD-ZIPIII genes contribute to vascular specification^*8,11,14*^, root and shoot apical meristem maintenance^*9,11,13*^, and the distinction of adaxial tissues in lateral organs^*9,10*^. These innovations parallel the increasing complexity of plant form and were instrumental to the enormous success of land plants^*1–3*^.

In keeping with a critical role in development, HD-ZIPIII activity is subject to intricate regulation. HD-ZIPIII transcripts are targeted by miR166, which restricts their accumulation via morphogen-like patterning properties^*15*^. Loss of this regulation conditions gain-of-function phenotypes impacting nearly all aspects of plant development^*10*^. At the protein level, HD-ZIPIII activity is modulated in part through interaction with their direct targets, the LITTLE ZIPPER (ZPR) family. ZPR proteins capture HD-ZIPIII TFs into heteromeric complexes lacking DNA-binding potential, creating a negative feedback loop that fine-tunes HD-ZIPIII activity^*16–18*^.

An additional layer of regulation is suggested by the fact that HD-ZIPIII proteins contain a START domain (**Fig. 1A**)^*19*^. START domains are members of the StARkin superfamily, which are present throughout the tree of life^*20,21*^. StARkin domains are characterized by an α/β helix-grip fold structure with a deep hydrophobic pocket that accommodates lipophilic ligands such as long-chain fatty acids, sterols, and isoprenoids^*22–24*^. Ligand binding induces stereotypical conformational changes that activate StARkin proteins through a diverse set of non-mutually exclusive regulatory mechanisms (reviewed in ^*21*^). For instance, StARkin domains can control protein turnover, homomeric and heteromeric complex stoichiometry, subcellular localization, and secondary structure stability^*25–34*^. The presence of a homeodomain and a START domain therefore sparked a long-standing hypothesis that the transcriptional activity of HD-ZIPIII proteins may be controlled by a lipid ligand, in a manner reminiscent of nuclear receptors in mammalian systems^*35–37*^. Remarkably, a role for the HD-ZIPIII START domain has not been established, despite the essential roles these TFs play in development^*9,10*^.

**Fig 1.**
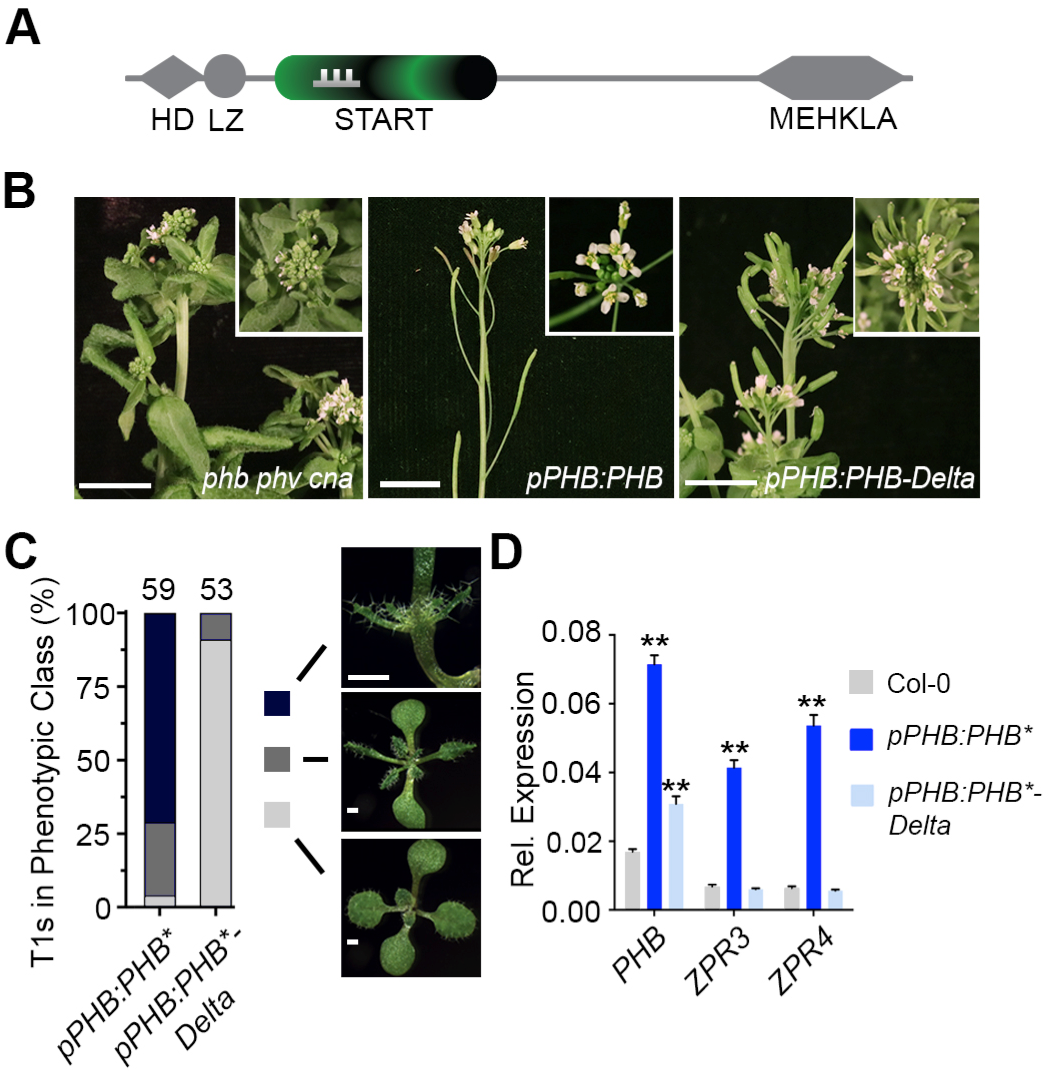
The START domain is required for full PHB function. **A**, General structure of HD-ZIPIII proteins. Note PHB-Delta retains the miR166 binding site (grey comb) contained within the START domain (green). **B**, The *pPHB:PHB* transgene complements the *phb phv cna* mutant phenotype while *pPHB:PHB-Delta* does not. **C**, Phenotypic scoring of primary transformants (*n* above bar) carrying miR166-insensitive (*) *pPHB:PHB* or *pPHB:PHB-Delta* constructs. Ectopic accumulation of PHB leads to severe (black) or intermediate (dark grey) gain-of-function phenotypes, whereas plants mis-accumulating PHB-Delta appear wild-type (light grey). **D**, Ectopic *PHB\** expression leads to strong upregulation of *ZPR3* and *ZPR4* targets. By contrast, *ZPR3* or *ZPR4* levels are indistinguishable from Col-0 in *pPHB:PHB*-Delta* lines, despite accumulation of this variant above endogenous *PHB* levels. Note: higher PHB transcript levels in *pPHB:PHB\** could reflect either a PHB autoactivation mechanism (proposed in ^*10*^) or the greater relative proportion of adaxialized tissues in *pPHB:PHB\** seedlings (versus *pPHB:PHB*-Delta*). *n* = 3 biological replicates. \*\**P* ≤ 0.01, Student’s *t*-test.

Potential insights may come from the evolutionarily related HD-ZIPIV family, which also diverged at least 725 million years ago^*1–3*^, and whose START domain is required for function^*29,32,33*^. Yeast expression analyses indicate that this START domain binds a broad spectrum of metabolites and increases TF stability^*29*^. Recent studies find similar promotive effects on protein stability in plants and identify subcellular localization as an additional HD-ZIPIV START regulatory mechanism controlling epidermal cell fate^*32,33*^. These effects are thought to be mediated by binding of epidermally-synthesized ceramides, which would reinforce their tissue-specific activity^*32,33*^. Given the long evolutionary divergence of HD-ZIPIII proteins^*1–3*^, their functions outside of the epidermis^*4–13*^, and the fact that HD-ZIPIII and HD-ZIPIV START domains are not interchangeable in yeast or plant assays^*29*^, the extent to which these observations apply to the HD-ZIPIII START domain is difficult to predict.

We show that addition of the HD-ZIPIII START domain potentiates transcriptional activity by promoting homodimerization and increasing transcriptional potency. Further, mutations predicted to affect ligand binding, or its downstream response, abolish DNA-binding competence, without overt effects on protein stability, subcellular localization, and interaction partners. Thus, the HD-ZIPIII START domain potentiates TF activity through regulatory mechanisms distinct from the evolutionarily related HD-ZIPIV START domain. These findings resolve a long-standing mystery in plant development and highlight the flexible and diverse regulatory potential coded within the ubiquitously distributed StARkin evolutionary module.

## Results

### The START domain is required for full PHB function

To assess START-dependent effects on PHB developmental function, we replaced this domain in the functional YFP-tagged *pPHB:PHB* reporter (Skopelitis et al., 2017) with the 21-nt miR166 recognition site found within the START domain coding sequence^*10*^ (*pPHB:PHB-Delta*). Whereas the *pPHB:PHB* transgene complements the *phb, phavoluta (phv*), and *corona (cna*) triple mutant phenotype, the *pPHB:PHB-Delta* construct fails to rescue (**Fig. 1B**), despite equivalent accumulation of *PHB* and *PHB- Delta* transcripts (**Fig. S1A**). We therefore next introduced a silent mutation into the 21-nt miR166 binding site, abolishing miR166 regulation of PHB and PHB-Delta (*pPHB:PHB*; pPHB:PHB*-Delta*). Loss of miR166 regulation generates a highly sensitive, dosage-dependent readout of HD-ZIPIII activity^*8,10,15*^, permitting detection of weak or subtle HD-ZIPIII function that might be missed in standard complementation assays. As expected^*38*^, over 90% of miR166-insensitive *pPHB:PHB\** primary transformants show PHB gain-of-function phenotypes (**Fig. 1C**). By contrast, *pPHB:PHB*-Delta* transformants are indistinguishable from wild-type plants (**Fig. 1C**), despite similar ectopic accumulation of protein (**Fig. S1B-E**). Further, transcript levels of the HD-ZIPIII direct targets *ZPR3* and *ZPR4* are strongly upregulated in *pPHB:PHB\** lines, but are indistinguishable from wild-type in *pPHB:PHB*-Delta* transformants (**Fig. 1D**). Thus, the START domain is required for PHB to fulfill its developmental function. Moreover, as PHB-Delta shows no obvious effects on protein stability or subcellular localization (**Fig. S1B-E**, and see below), the HD-ZIPIII START domain employs regulatory mechanisms distinct from those of the evolutionary related HD-ZIPIV START domain.

### The START domain promotes PHB dimerization

One frequently used StARkin regulatory mechanism is modulation of homomeric stoichiometry^*24,34,39–43*^. To test whether the START domain impacts PHB homodimerization, we used single-molecule pull down (SiMPull), which we previously adapted for plant systems^*18*^. We first determined the maturation frequency of the monomeric citrine YFP variant in *Arabidopsis* to calibrate the frequency of two-step photobleaching events into a quantitative assessment of homomeric stoichiometry^*18,44–46*^ (**Figs. 2A** and **S2)**. Subsequent analyses of over 3300 protein complexes showed that ~80% of wild-type PHB proteins are present as dimers (**Fig. 2B**). This frequency resembles that of strongly-homodimeric proteins in animal systems^*44,46*^, and predicts PHB functions primarily as a homodimer. By contrast, PHB-Delta has a dimerization frequency of 15% – about twice the frequency with which two-step photobleaching events are observed by chance (**Fig. 2B** and **Fig. S2B**). Thus, the PHB START domain promotes either the formation or the maintenance of PHB homodimers (**Fig. 2B**). As HD-ZIPIII proteins require dimerization to bind DNA^*47*^, this effect on homodimerization provides a potential explanation for PHB-Delta failing to activate the normal PHB developmental program (**Figs. 1B-D**).

**Fig 2.**
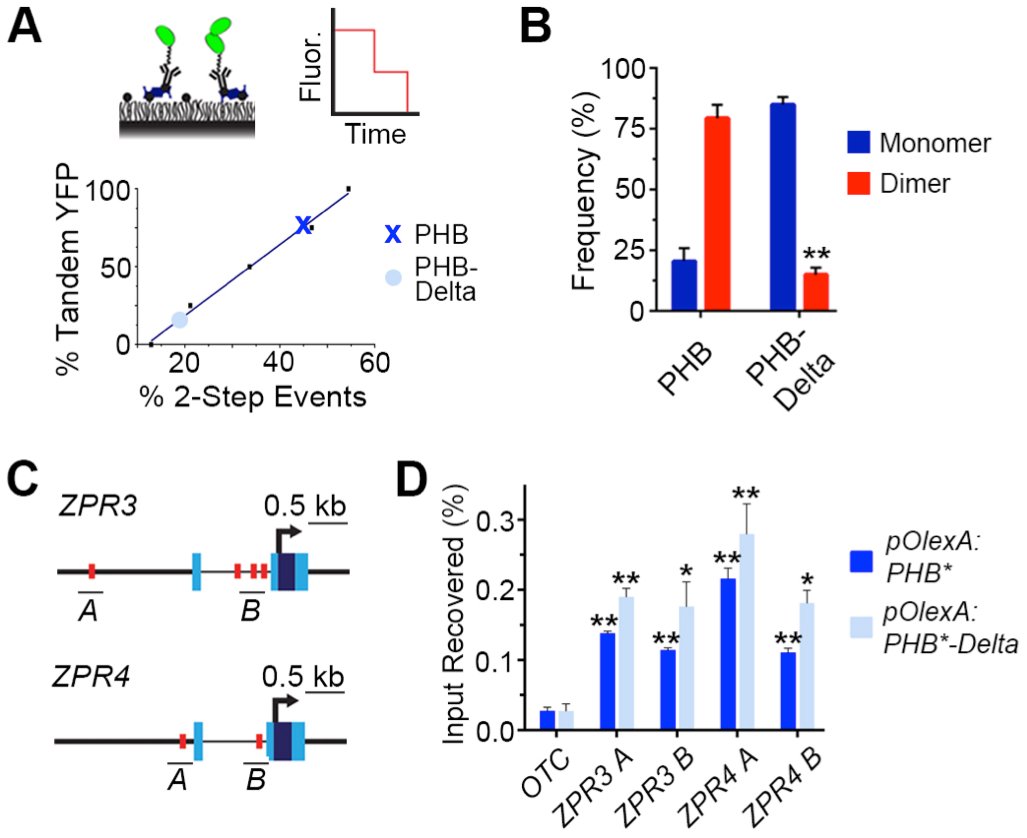
The START domain affects PHB homodimerization. **A**, SiMPull calibration curve (bottom graph) for monomeric vs dimeric YFP (top left cartoon) translates percentage of two-step photobleaching events (simulated in top right cartoon) to frequency of dimers in the population. PHB (**X**) and PHB-Delta (•) show two-step photobleaching events of 47% and 20%, respectively. **B**, Percentages of two-step photobleaching events translate to dimerization frequencies of ~80% for PHB and ~15% for PHB-Delta. **C**, Schematic representations of *ZPR3* and *ZPR4* showing HD-ZIPIII binding sites identified by FIMO (red ovals), ChIP amplicons (black bars), transcription start sites (arrow), untranslated regions (light-blue boxes), and exons (dark-blue boxes). **D**, PHB and PHB-Delta occupy multiple sites in the regulatory regions of *ZPR3* and *ZPR4* and are significantly enriched over the *ORNITHINE TRANSCARBAMILASE (OTC*) negative control locus. *n* = 3 biological replicates. \**P* ≤ 0.05, \*\**P* ≤ 0.01, Student’s *t*-test.

### The START domain enhances PHB transcriptional potency

One consequence of poor dimerization of PHB-Delta is that it may obscure possible additional contributions from the START domain to HD-ZIPIII TF function. We therefore turned to a short-term estradiol-inducible overexpression system to increase the total number of PHB-Delta dimers available for molecular assays of TF activity. Importantly, estradiol induction does not change the subcellular localization or dimerization frequencies of PHB or PHB-Delta seen at native expression levels (**Figs. S3A-S3D**). Both variants can also be induced to similar levels and show comparable protein stability (**Figs. S3A** and **S3E**), making this approach suitable for identifying additional START-dependent effects.

Using short-term estradiol-inductions, we first tested whether PHB proteins lacking the START domain are capable of binding to DNA using chromatin immunoprecipitation (ChIP). PHB occupies multiple palindromic HD-ZIPIII binding sites^*47*^ in the regulatory regions of its *ZPR3* and *ZPR4* direct targets (**Figs. 2C** and **2D**). Similarly, enrichment at these sites was also detected for PHB-Delta (**Figs. 2C** and **2D**), indicating this variant retains the capacity to bind to DNA.

Given this outcome, we next gauged the effect of the START domain on transcriptional potency. Here, the short-term estradiol-induction system has the added benefit that it provides a direct quantitative readout of transcriptional activity while avoiding confounding consequences of morphological changes and regulatory feedback^*16–17*^. As expected, *ZPR3* and *ZPR4* transcript levels are strongly upregulated upon induction of PHB (**Fig. 3A**). *ZPR3* and *ZPR4* targets are also upregulated upon induction of PHB-Delta (**Fig. 3A**). Intriguingly, these transcripts are upregulated to between one-half and one-third the levels seen for PHB, despite equivalent induction of *PHB* and *PHB-Delta* (**Fig. 3A**). As PHB variants are not differentially stable (**Figs. S1B-E** and **S3E**), these data suggest deletion of the START domain, in addition to reducing the frequency of dimers, significantly reduces their transcriptional potency. This idea is supported by two orthogonal lines of evidence. First, virtually identical results were obtained using cotransfection assays in *Nicotiana benthamiana*, which show PHB-Delta fails to fully activate both endogenous *ZPR* targets and a *pZPR3:3x-NLS-RFP* reporter (**Fig. S4**). Second, the phenotypic severity of transformants constitutively overexpressing PHB-Delta (*p35S:PHB*-Delta*) is markedly lower than the *p35S:PHB\** control (**Fig. S5A**). *ZPR3* and *ZPR4* transcripts in *p35S:PHB*-Delta* lines also accumulate to between one-half and one-third the levels seen in *p35S:PHB\** seedlings, despite equivalent accumulation of *PHB* and *PHB-Delta* transcripts (**Fig. S5B**). These complementary assays support augmenting of transcriptional potency as an additional HD-ZIPIII START regulatory mechanism.

**Fig 3.**
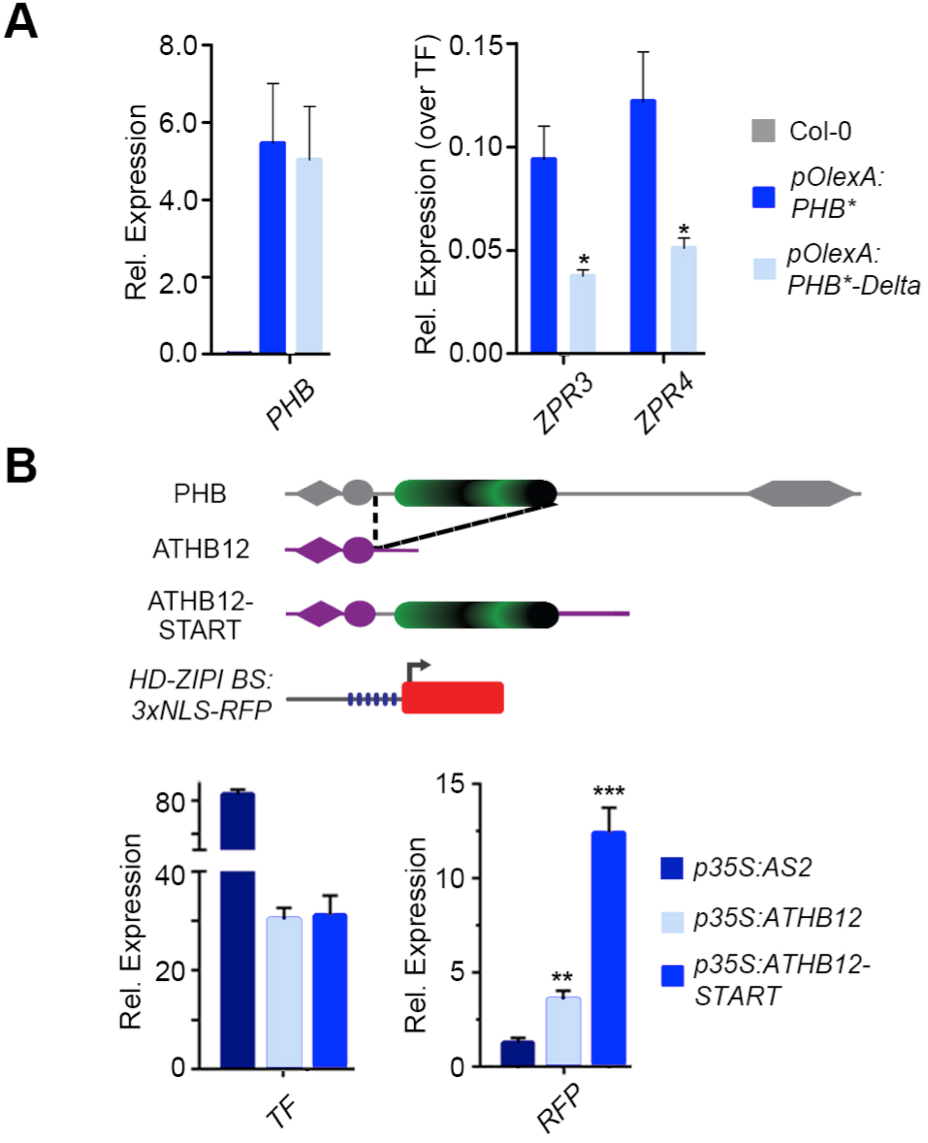
The START domain enhances PHB transcriptional potency and this property is transferable to heterologous TFs. **A**, Relative *ZPR3* and *ZPR4* transcript levels in 24 hrs estradiol-induced *pOlexA:PHB\** and *pOlexA:PHB*-Delta* seedlings indicate PHB-Delta is a less-potent transcriptional activator than PHB. **B**, Schematic representation of domain-capture-mimicry constructs detailing insertion of the PHB START domain (green) into ATHB12 (purple) and the design of the *HD-ZIPI BS:3xNLS-RFP* target reporter (top). qRT-PCR following co-transfection with the *HD-ZIPI BS:3xNLS-RFP* target reporter (bottom), shows fusion of the wild-type PHB START domain augments ATHB12 transcriptional potency by a factor of three (bottom). AS2 is included as negative control. *n* = 3 biological replicates. \**P* ≤ 0.05, \*\**P* ≤ 0.01, Student’s *t*-test.

### START regulatory properties are transferable onto heterologous TFs

The idea that the PHB START domain potentiates TF activity and is not strictly required for TF identity *per se* is in line with principles of TF evolution by domain capture^*48–51*^. Relevant to this, plant genomes encode HD-ZIP TFs that lack a START domain^*1–3*^. These related HD-ZIP members provide a unique opportunity to definitively test how addition of a START domain impacts TF output. We therefore inserted the PHB START domain downstream of the homeodomain and leucine zipper motifs of the HD-ZIPI family member ATHB12, partially recapitulating HD-ZIPIII architecture (ATHB12-START). ATHB12 is a known activator of transcription, and is sufficient to drive expression of reporters placed downstream of multimeric HD-ZIPI binding sites^*52*^ (**Fig. 3B**). Remarkably, addition of the HD-ZIPIII START domain increases ATHB12 transcription activity by a factor of three over unmodified ATHB12 (**Fig. 3B**). This effect on transcriptional potency parallels effects seen for PHB and PHB-Delta in *Arabidopsis* as well as equivalent co-transfection assays in *N. benthamiana* (**Figs. 3A** and **S4**). Thus, the START domain is necessary for HD-ZIPIII dimers to achieve full transcriptional potency, and its addition is sufficient to confer this increase in potency onto heterologous TFs.

### The START domain does not affect heteromeric stoichiometry

At a molecular level, StARkin domains undergo stereotypical conformational changes in response to ligand binding^*22,23*^. In the *apo* form, StARkin domains reveal an open ligand-binding pocket, and upon ligand interaction the pocket is sealed via conformational changes. In addition to changes in tertiary structure, this creates a new interaction surface that, for a subset of StARkin proteins, mediates recruitment of additional protein partners to exert StARkin-dependent regulatory effects^*24,39–43*^. To test this possibility, we performed quantitative mass spectrometry on proteins that co-immunoprecipitate with PHB (IP-MS; **Fig. S6**). Among the proteins significantly enriched across five PHB IP-MS replicates were multiple members of the BRAHMA-containing SWI/SNF chromatin-remodeling complex. These data point to specificity of the IP-MS and suggest HD-ZIPIII proteins facilitate transcription via direct modification of chromatin. Further supporting specificity of the IP-MS, the known HD-ZIPIII interacting partners ZPR1 and ZPR3^*16–18*^ also co-immunoprecipitate with PHB. Finally, significant enrichments were detected for the lipid-binding proteins OLEO1, OLEO5, DRP1C, DRP1E, and ANNATD4, which is of potential interest given the lipid-binding nature of StARkin domains (**Fig. S6** and **Table S1**). Proteinprotein interactions are, however, not mediated via the START domain as IP-MS shows that PHB-Delta binds the same interaction partners (**Table S2**), and SiMPull co-localization analyses confirm that interaction with ZPR3 is unchanged (**Fig. S7**). These findings thus argue against START-mediated regulation of HD-ZIPIII complex stoichiometry at the level of interaction partners.

Taken together, complementary phenotypic and molecular analyses reveal the START domain is required for PHB developmental function, but unlike its homolog in the HD-ZIPIV TFs, does not impact protein stability or subcellular localization. Instead, the presence of a START domain in PHB potentiates TF activity by promoting homodimerization and by increasing transcriptional potency. These effects seem to be mediated by interactions between domains of PHB, as the START domain does not appear to determine its spectrum of interaction partners.

### Mutating conserved START residues abolishes PHB DNA-binding competence

Following the idea that, like other StARkin domains^*38,41*^, the HD-ZIPIII START domain influences intramolecular domain-to-domain interactions, mutations perturbing this property may exert effects on TF function distinct from those observed after full deletion of the domain. To assess this possibility, we first performed homology modeling of the PHB START domain using I-TASSER and AlphaFold2 to identify conserved functional residues to target via mutagenesis. Both algorithms indicate the START domain is distinct from sterol- or isoprenoid-binding StARkin domains such as the ABA receptor^*43*^, and instead resembles mammalian phosphatidylcholine transfer protein^*23*^ (PC-TP; **Fig. 4A**). START domains like PC-TP contain several highly conserved residues that mediate the ligand-directed conformational change^*19,22,23,53*^. Accordingly, two short amino acid stretches (RDFTWLR and RAEMK), centered around three such arginine residues^*19,22,23,53*^, were selected for mutagenesis (SDmut; **Figs. 4A** and **S8**). Importantly, these mutations are predicted to minimally affect protein folding (RMSD 0.291; **Fig. 4A**), and circular dichroism confirmed that wild-type, SDmut, and PC-TP START domains purified from *E. coli* adopt virtually identical secondary structures (**Fig. 4B**).

**Fig 4.**
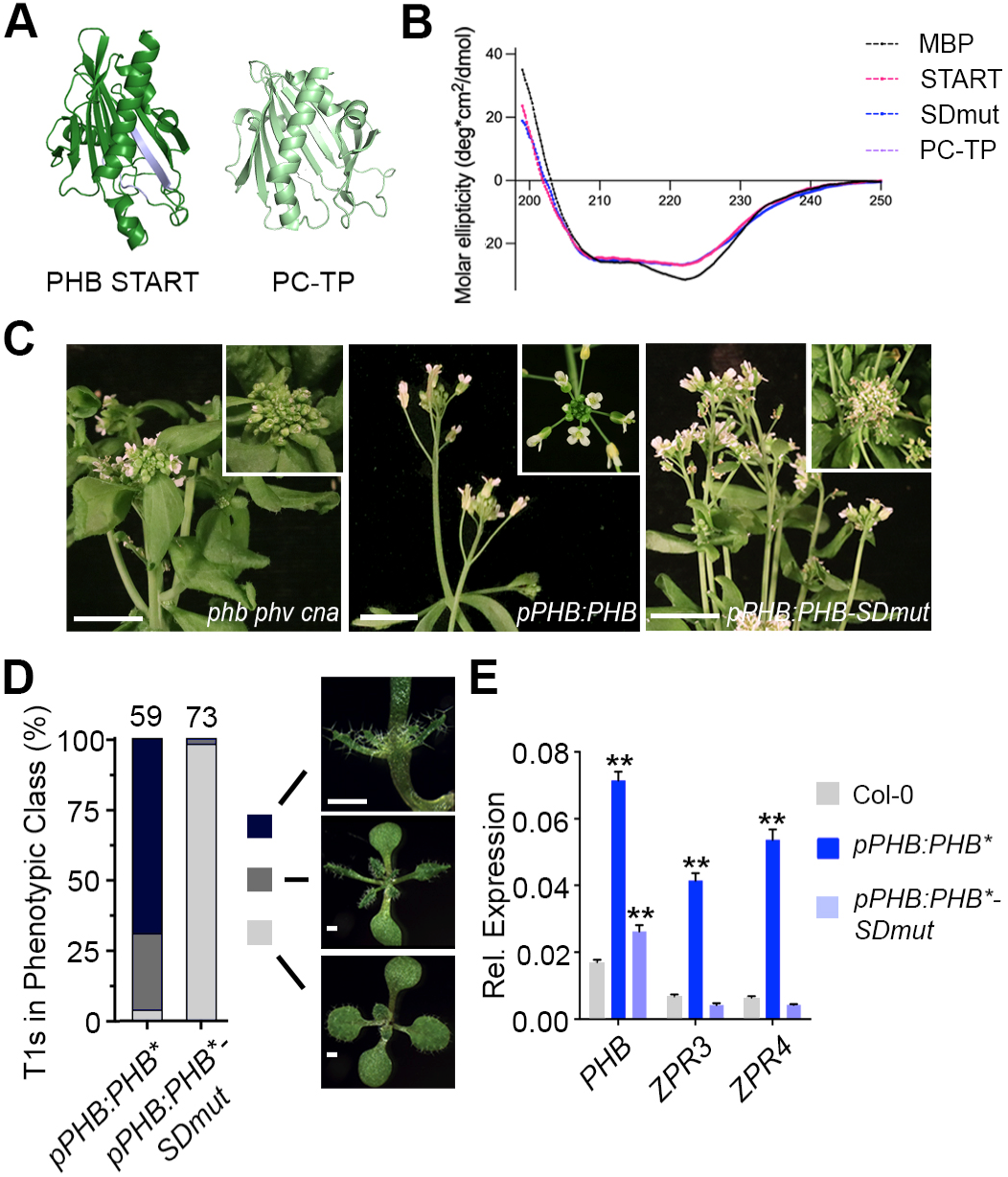
Mutating the START domain perturbs PHB developmental function. **A**, Homology-guided modeling predicts the PHB START domain (left) closely resembles PC-TP (right). Positions of amino acids mutated in PHB-SDmut are shown in lavender. **B**, Circular dichroism of MBP (negative control), MBP-START, MBP-SDmut, and MBP-PC-TP shows all three recombinant START proteins have virtually identical secondary structures. **C**, The *pPHB:PHB* transgene complements the *phb phv cna* mutant while *pPHB:PHB-SDmut* does not. **D**, Phenotypic scoring of primary transformants (*n* above bar) carrying miR166-insensitive (*) *pPHB:PHB* or *pPHB:PHB-SDmut* constructs. Ectopic accumulation of PHB leads to severe (black) or intermediate (dark grey) gain-of-function phenotypes, whereas plants mis-accumulating PHB-SDmut appear wild-type (light grey). **E**, Ectopic *PHB\** expression leads to strong upregulation of *ZPR3* and *ZPR4* targets. By contrast, *ZPR3* or *ZPR4* levels are indistinguishable from Col-0 in *pPHB:PHB*-SDmut* seedlings, despite accumulation of this variant above endogenous *PHB* levels. *n* = 3 biological replicates. \*\**P* ≤ 0.01, Student’s *t*-test. Note: PHB, PHB-SDmut, and PHB-Delta data were collected simultaneously. PHB data from **Fig. 1** are replotted for clarity.

Identical mutations were then introduced into the functional YFP-tagged *pPHB:PHB* reporter and its miR166-insensitive counterpart *pPHB:PHB\**, creating *pPHB:PHB-SDmut* and *pPHB:PHB*-SDmut*, respectively. The *pPHB:PHB-SDmut* transgene failed to complement the *phb phv cna* mutant phenotype (**Fig. 4C**), suggesting the selected residues are indeed required for PHB function. Supporting this, *pPHB:PHB*-SDmut* primary transformants did not show phenotypes in the more-sensitive dosedependent, gain-of-function assay (**Fig. 4D**), and *ZPR3* and *ZPR4* transcript levels in these lines were indistinguishable from the wild-type (**Fig. 4E**). These effects are again not explained by changes in protein stability, sub-cellular localization, or interacting partners, as these properties are comparable for PHB, PHB-Delta, and PHB-SDmut (**Figs. S1B-E, S3, S7**, and **Table S2**). By contrast, SiMPull showed that homodimeric PHB-SDmut complexes are present at ~40% frequency in the population (**Fig. 5A**), approximately twice that seen for PHB-Delta but half that seen for wild-type PHB (**Fig. 2B**).

**Fig. 5.**
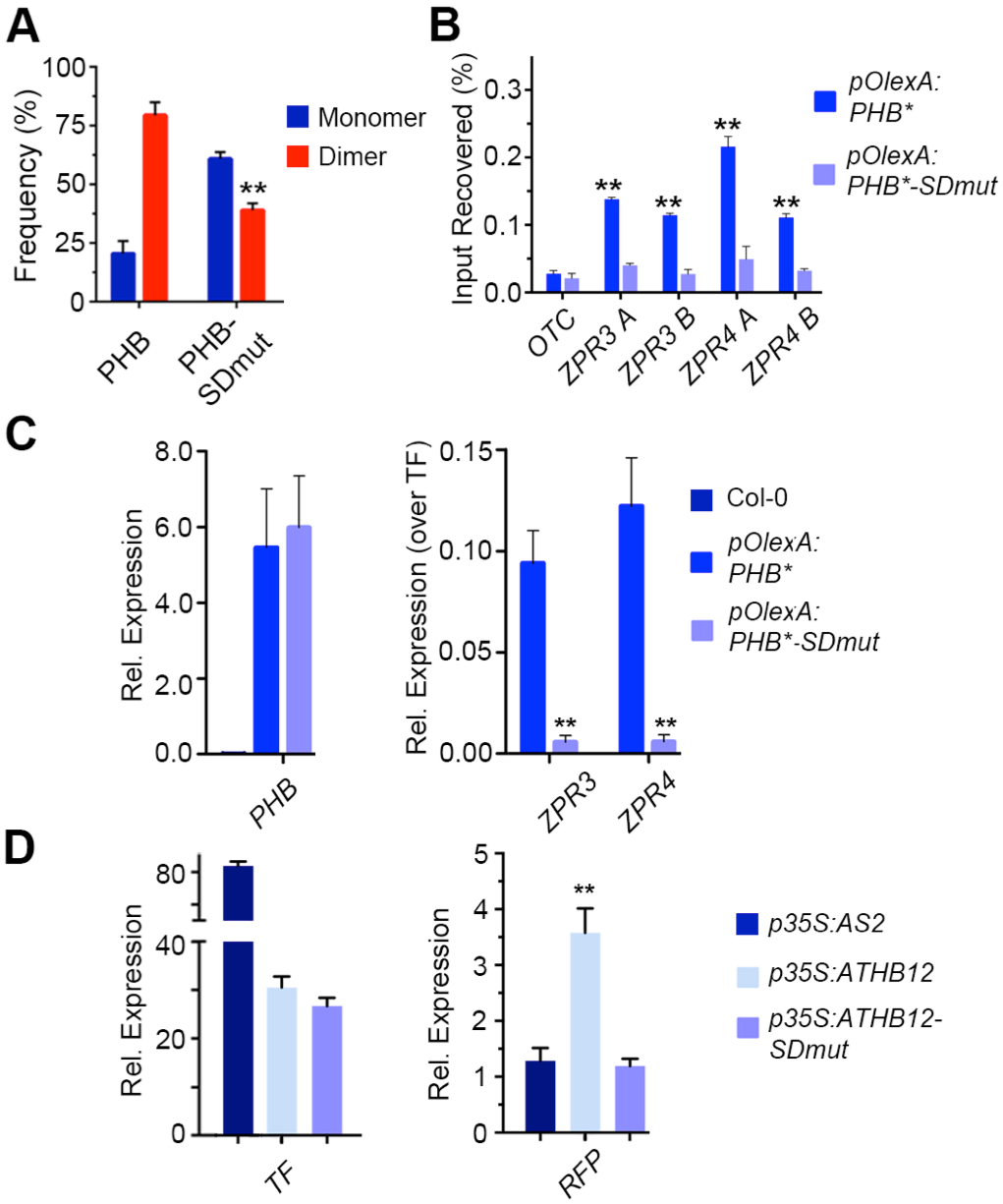
Mutating PHB START reduces homodimerization and abolishes DNA-binding competence. **A**, Percentages of two-step photobleaching events from SiMPull translate to dimerization frequencies of ~80% for PHB and ~40% for PHB-SDmut. **B**, PHB-SDmut does not bind *ZPR3* or *ZPR4* regulatory regions occupied by PHB. **C**, Unlike PHB, PHB-SDmut cannot activate *ZPR3* or *ZPR4* targets in 24h estradiol-induction experiments. **D**, Domain-capture-mimicry experiments demonstrate fusion of SDmut START to ATHB12 (ATHB-SDmut) renders ATHB12 non-functional, as RFP reporter levels are indistinguishable from the AS2 negative control. *n* = 3 biological replicates. \*\**P* ≤ 0.01, Student’s *t*-test. Note: PHB, PHB-SDmut, and PHB-Delta data were collected simultaneously. PHB data from **Figs. 2** and **3** are replotted for clarity.

We then tested whether PHB-SDmut dimers retain DNA-binding capability. Interestingly, no enrichment of PHB-SDmut was detected in the regulatory regions of the *ZPR3* or *ZPR4* loci using ChIP (**Fig. 5B**), indicating these mutations abolish PHB DNA-binding competence. Consistent with this, PHB-SDmut fails to activate *ZPR3* and *ZPR4* in short-term estradiol-induction assays in *Arabidopsis* as well as in co-transfection assays in *N. benthamiana* (**Figs. 5C** and **S4**). Moreover, addition of this non-functional START domain to ATHB12 abolishes its transcriptional activity (ATHB12-SDmut; **Fig. 5D**). Thus, mutating the START domain, and deleting the START domain entirely, leads to proteins with distinct biochemical properties: PHB-SDmut dimerizes relatively well but is unable to bind DNA, whereas PHB-Delta rarely dimerizes but those dimers that do form retain DNA-binding competence and have reduced transcriptional potency.

### The PHB START domain binds PC and mutating predicted PC-binding residues abolishes DNA-binding

Structural modeling and mutational analyses propose the PHB START domain may be controlled by phospholipid ligands (**Figs. 4A**, **5**, and **S8**). We therefore tested whether the PHB START domain interacts with a set of ligands similar to those bound by PC-TP. Depending on context, these include members of the PC, phosphatidylethanolamine (PE), and phosphatidylglycerol (PG) phospholipid classes^*54–56*^. To this end, recombinant PHB START domain protein was incubated with liposomes generated from *Arabidopsis* total lipid extracts, re-purified by affinity chromatography, and then subjected to LC-MS at two independent lipidomics centers. Prior to incubation with plant-derived liposomes, recombinant PHB START protein co-purified with bacterial PE and PG species (**Figs. S9A, S9B, Table S3**). Binding of these fortuitous “ligands” has also been reported for other recombinant PC-binding proteins including PC-TP^*54*^, steroidogenic factor 1 (SF-1)^*55*^, and liver receptor homolog-1 (LRH-1)^*56*^. After incubation with plant-derived liposomes (**Fig. S9C**), additional lipids were significantly enriched across five PHB START LC-MS replicates (**Tables S4-S6**). These include five species of PC, two of which are preferred ligands for PC-TP^*54*^ (**Fig. 6A**). Further, membrane-overlay assays indicate this binding is not occurring at the surface of the START domain (**Fig. S9D**), hinting at further parallels between the HD-ZIPIII and PC-TP START domains.

**Fig. 6.**
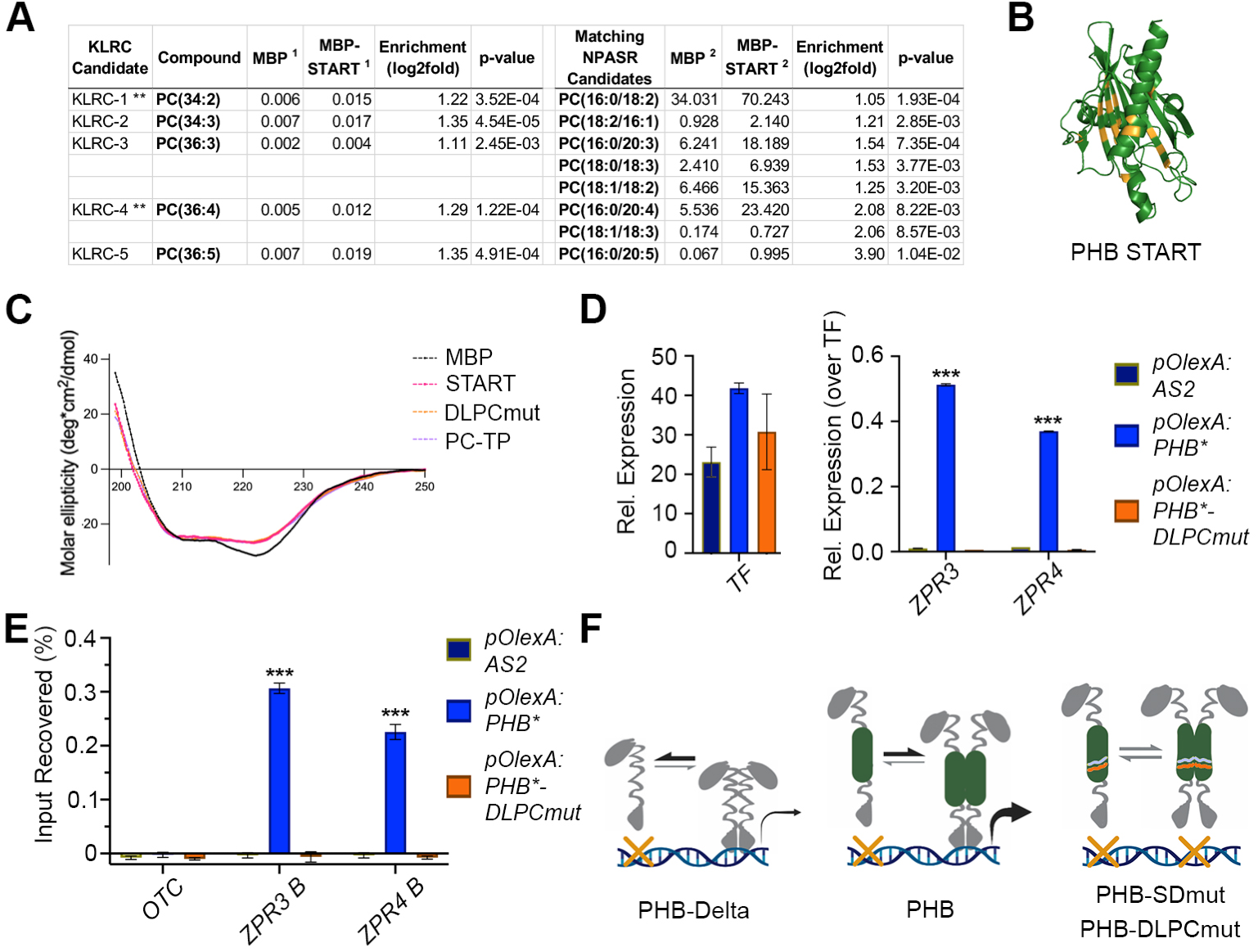
PHB START domain binds PC and mutating predicted PC-binding residues abolishes DNA-binding. **A**, LC-MS analysis of lipids bound by recombinant MBP and MBP-START proteins after liposome incubation and repurification. Several species of PC are significantly enriched in MBP-START. Raw signal intensities are reported, ** indicates species preferentially bound by PC-TP^*31*^. **B**, Homology modeling of DLPCmut START variant showing position of mutations (orange). **C**, Circular dichroism of MBP, MBP-START, MBP-DLPCmut, and MBP-PC-TP showing all three recombinant START proteins have virtually identical secondary structures. **D**, Unlike PHB, PHB-DLPCmut cannot activate *ZPR3* or *ZPR4* targets in 24h estradiol-induction experiments. **E**, PHB-DLPCmut does not bind *ZPR3* or *ZPR4* regulatory regions occupied by PHB. **F**, Without a START domain, HD-ZIPIII TFs make fewer and less transcriptionally potent dimers (left), while mutations predicted to eliminate START activity abolish DNA binding (right; START in green; mutations in lavender and orange). These data propose a model in which a functional START domain promotes HD-ZIPIII dimerization and transcriptional potency, but requires a ligand for these TFs to bind DNA (middle). *n* = 3 or 4 biological replicates. \*\*\**P* ≤ 0.001, Student’s *t*-test.

We therefore created a new PHB START variant with mutations in seventeen residues analogous to those contacting PC within the PC-TP binding pocket^*23*^ (DLPCmut; **Figs. S8, S9E**). The substitutions chosen have minimal effects on overall structure (RMSD 0.372; **Figs. 6B**), and include six amino acids with bulkier side chains that partially occlude the ligand binding pocket (**Figs. S9F-G**). Importantly, circular dichroism confirmed that purified wild-type, DLPCmut, and PC-TP START domains have nearly indistinguishable secondary structures (**Fig. 6C**). PHB proteins with the DLPCmut START domain (PHB-DLPCmut) are nuclear-localized in plants but fail to activate *ZPR3* and *ZPR4* (**Fig. 6D; Fig. S9H**). Moreover, no enrichment of PHB-DLPCmut was detected at *ZPR3* using ChIP assays (**Fig. 6E**). Taken together, these data propose species of PC as promising candidate ligands for the HD-ZIPIII START domain and show that mutations in amino acids predicted to affect either ligand-binding or ligand-mediated conformational change result in similar loss of HD-ZIPIII DNA-binding competence (**Figs. 4B** and **6E**).

## Discussion

HD-ZIPIII TFs are principal regulators of key developmental innovations throughout plant evolution. Although molecularly cloned over twenty years ago^*10*^, the contribution of their START domain remained elusive. We show that the START domain promotes TF homodimerization and increases transcriptional potency. These effects are mediated solely through intramolecular protein changes and can be ported onto other TFs. These findings are particularly intriguing when considered through the lens of TF evolution by domain capture. Basic HD-ZIP proteins and minimal START proteins resembling PC-TP are both present in unicellular algae^*2,3*^, whereas HD-ZIPIII architecture arose after the divergence of *Chlorophyta*, but before the emergence of *Chlorokybus atmophyticus* over 725 million years ago^*57*^. Capture of a START domain by HD-ZIPIII antecedents may thus have augmented their transcriptional output. Presumably this also placed their activity under control of a ligand such as PC (**Fig. 6A**). This supposition is consistent with our data. We find that perturbing START domain residues predicted to affect either ligand-binding or downstream conformational change^*19,22,23,53,54*^ abolishes DNA-binding competence (**Figs. 5B** and **6E**). One potential explanation is that the *apo* form of the START domain holds PHB in a non-DNA-binding conformation. Binding of its ligand, or deletion of the START domain entirely, would relieve this inhibition and permit DNA binding. Taken together, these data, as well as the known properties of StARkin domains^*21–24,34,58*^, present a model in which the START domain potentiates HD-ZIPIII TF activity, and uses ligand-induced conformational change to render HD-ZIPIII dimers competent to bind DNA (**Fig. 6F**).

Given this model, it is intriguing to speculate whether acquisition of a START domain enabled HD-ZIPIII TFs to more effectively integrate signaling inputs into their gene networks, a critical feature of multicellularity^*59*^. In addition, as phospholipid accumulation is spatially regulated^*60,61*^, acquisition of the START domain could have allowed HD-ZIPIII activity to become patterned across groups of cells. This would have clear developmental implications as transcripts of ancestral HD-ZIPIII genes are not targeted by small RNAs of the miR166 family^*62*^. Such a regulatory paradigm draws parallels between HD-ZIPIII TFs and mammalian nuclear receptors^*37,63*^, however future experiments are needed to determine whether START ligands play structural roles or indeed contribute to the intricate spatiotemporal regulation of HD-ZIPIII activity. Either way, the properties of the START domain described here suggest a compelling basis for the emergence of HD-ZIPIII TFs as key drivers of plant morphogenic evolution.

The START domain of the closely related HD-ZIPIV family is similarly required for function, and impacts subcellular localization and protein stability, possibly through binding of epidermally-synthesized ceramides^*29,32,33*^. The regulatory properties conferred by the HD-ZIPIII START domain are thus distinct, despite their evolutionary relationship^*19,35*^. Further, the modulation of DNA-binding competence identified here represents a new type of StARkin-directed regulatory mechanism. Our findings, in addition to resolving a long-standing mystery in plant development, highlight the flexible and diverse regulatory potential coded within this widely distributed evolutionary module^*21*^.

## Methods

### Plant materials and growth conditions

*Arabidopsis thaliana* (Col-0 ecotype) seedlings and *Nicotiana benthamiana* (tobacco) plants were grown at 22°C under long-day conditions, on soil or 1% agarose plates containing Murashige and Skoog medium supplemented with 1% sucrose. Inductions were performed by spraying 10-day-old seedlings with 20 uM B-estradiol in 1% DMSO supplemented with 0.005% Silwet.

### Molecular biology and plant transformations

*pPHB:PHB-YFP* and *pPHB:PHB*-YFP* constructs have been described previously^*15*^. Using these templates, *pPHB:PHB-Delta-YFP* and *pPHB:PHB*-Delta-YFP* were constructed via Gibson assembly (NEB), replacing the 642bp START domain (496-1137bp from the start codon) with a sensitive (GGGATGAAGCCTGG**T**CCGGAT) or an insensitive version (GGGATGAAGCCTGG**A**CCGGAT) of the 21nt miR166-recognition site^*15*^. To create *pPHB:PHB- SDmut-YFP*, we first synthesized a mutated variant of the START domain (Mr. Gene), which replaced amino acids RDFTWLR with GAVVGAG and amino acids RAEMK with VAAGV by including the following nucleotide substitutions: CGTGACTTTTGGACGCTGAGA at position 841-861 bp from the start codon to GGTGCCGTCGTAGGAGCAGGC, and AGAGCTGAAATGAAA at position 961-975 bp from the start codon to GTGGCGGCCGGCGTC. miR166-insensitive *pPHB:PHB*-SDmut-YFP* was then constructed via site-directed mutagenesis (Stratagene), copying the mutations in *pPHB:PHB*-YFP*^*15*^. All constructs were shuttled into the pB7GW binary vector via Gateway LR reactions (VIB Ghent; Invitrogen).

To facilitate subsequent stable and inducible overexpression in *Arabidopsis*, and transient expression in tobacco, *PHB*-YFP*, *PHB*-SDmut-YFP*, and *PHB*-Delta-YFP* cDNAs were reamplified and cloned into pCR8-GW via Gibson assembly (Invitrogen; NEB). Gateway LR reactions then shuttled each cDNA into pEARLYGATE100 (stable overexpression), modified pMDC7 with the UBQ10 promoter in place of G10-90 (inducible overexpression), or p502Ω (transient expression; VIB Ghent). *PHB-DLPCmut*-YFP* cDNA was generated by replacing the wildtype START domain with a gene-synthesized mutant variant (GeneArt) using Gibson cloning (ThermoFisher). A Gateway LR reaction then shuttled *PHB-DLPCmut- YFP* into pMDC7. *ATHB12* was amplified from cDNA generated from seedlings, then cloned into pCR8-GW using Gibson (NEB). Fusions between *ATHB12* and the START domain variants were created by inserting a fragment of *PHB\** or *PHB-SDmut* (376-1137bp from the start codon) into *ATHB12* (at position 366 from the start codon) using Gibson assembly (Invitrogen; NEB). Gateway LR reactions then shuttled these cDNAs into p502Ω (Invitrogen; VIB Ghent).

The *pZPR3:3xNLS-RFP* reporter was created by first cloning the 3.2kb region upstream of the *ZPR3* start codon into pCR8-GW (Invitrogen; NEB), followed by an LR reaction using a modified pGREEN binary vector with a Gateway cassette upstream of *3xNLS-RFP* (Invitrogen). The *HD-ZIPI BS:3xNLS-RFP* reporter was created by first synthesizing a construct comprised of six copies of the HD-ZIPI binding site (CAATTATTG) followed by a minimal 35S enhancer, flanked by attL1/attL2 sites (Mr. Gene). A 10-nt spacer (CATTTCAAGA) was inserted between each binding site to minimize potential stearic hindrance. Finally, an LR reaction was used to shuttle these multimerized binding sites into the pGREEN binary described above (Invitrogen). Cloning primer sequences for all constructs are listed in **Table S7**. Synthesized sequences are in **Table S8**.

Transient transfection of tobacco was performed using syringe-mediated infiltration^*64*^. In brief, overnight cultures of *Agrobacterium tumefaciens* were centrifuged, resuspended in 2 to 5mL of room temperature infiltration medium (10 mM MgCl2, 10 mM MES KOH, pH 5.6, 150 mM acetosyringone [Sigma Aldrich], and 1% DMSO), diluted to a working optical density of 1, and infiltrated into third and fourth leaves of 3- to 4-week-old tobacco plants.

### Homology modeling and sequence alignment

The PHB START domain was modeled with I-TASSER (https://zhanggroup.org/I-TASSER/) and AlphaFold2 (https://colab.research.google.com/github/sokrypton/ColabFold/blob/main/AlphaFold2.ipynb) using default parameters. Sequences were aligned using the ClustalW algorithm in the MEGA X software (v. 10.1.7; https://www.megasoftware.net/). The crystal structure of PC-TP (1ln1) was the top template used for threading in I-TASSER. Models were visualized in Pymol.

### ChIP and qRT-PCR assays

ChIP assays were performed with 11-day-old estradiol-induced seedlings as previously described^*65*^, using IgG (abcam ab46540) or anti-GFP (abcam ab290) antibodies. ChIP and input DNA samples were assayed by qPCR using iQ SYBR Green Supermix (Bio-Rad). *ZPR3* and *ZPR4* regulatory regions assayed in ChIP were selected based on the presence of HD-ZIPIII binding sites predicted by FIMO (https://meme-suite.org/meme/tools/fimo). All experiments were performed at least three independent times. PCR was performed in duplicate, and enrichments calculated relative to input. Student’s t test was used to calculate statistical significance.

Total RNA was extracted from seedlings or infiltrated tobacco leaves using Trizol reagent (Gibco BRL). One microgram of RNA was primed with oligo (dT) and reverse transcribed using the SuperScript III first-strand synthesis kit (Invitrogen). Relative quantification values were calculated based on at least three biological replicates, with ΔCt of ACT2 or B-tubulin serving as normalization controls in *Arabidopsis* or tobacco, respectively. Wild-type or uninduced values were set to one and PHB variant values either plotted directly or after further normalization to PHB variant levels. Student’s t test was used to calculate statistical significance. ChIP and qRT-PCR primer sequences are listed in **Table S9**.

### Plant imaging

Brightfield images of *Arabidopsis* seedlings were captured using an SMZ1500 dissecting microscope with NIS Element software (Nikon). Fluorescent images of *Arabidopsis* seedlings and infiltrated tobacco leaves were obtained using the same microscope with the P-FLA2 epi-fluorescent attachment. Heart-stage embryos were dissected, stained with Fluorescent Brightener 28 (Sigma Aldrich), and then imaged using an LSM780 confocal microscope (Zeiss).

### Single-molecule imaging and stoichiometric analyses

A detailed protocol of SiMPull with plant tissue has been published^*18*^, however the amount of input tissue can vary between experiments. Here, lysates for SiMPull were prepared from flash frozen tissue comprised of five-to-six 10-day-old *Arabidopsis* seedlings or 1-2 cm^2^ pieces of infiltrated tobacco leaves. Note: SiMPull experiments in *Arabidopsis* were conducted using seedlings expressing PHB variants under native regulatory elements (**Figs. 2A-B**) as well as an estradiol-inducible promoter (**Fig. S3D**). A minimum of three independent biological replicates were performed for each SiMPull experiment. To enable calculation of PHB variant dimerization frequencies, as well as the maturation probability of citrine YFP in *Arabidopsis, 2×35S:monoYFP* and *2×35S:tdYFP* constructs (described in ^*18*^) were stably transformed and at least four independent lines analyzed.

SiMPull was first performed with 1:150 dilutions of lysates from *2×35S:monoYFP* or *2×35S:tdYFP* seedlings, mixed at ratios described in **Fig. S2**. Frequencies of two-step photobleaching events were then scored and used to construct a standard calibration curve (y = 2.28x - 27; R^2^ = 0.992). For stoichiometric analyses of PHB variants, lysates from native promoter lines or 24 hrs estradiol-induced lines were diluted 1:1 or 1:100, respectively, then subject to SiMPull and photobleaching counts. Frequencies of two-step photobleaching events for each genotype were translated into dimerization frequencies via the above calibration curve. Maturation probability was predicted using binomial probability modeling (described in ^*18*^). Co-localization data for ZPR3-mCherry with AS2-YFP, PHB-YFP, PHB-SDmut-YFP, or PHB-Delta-YFP were collected and analyzed as described previously^*18*^.

### Quantification of PHB variant induction with estradiol

Ten-to-twelve 10-day-old seedlings were induced for 24 hrs with estradiol, flash frozen, then ground in 500 μl freshly prepared lysis buffer (25 mM Tris HCl pH 8, 150 mM NaCl, 1% SDS, 1x cOmplete ULTRA protease inhibitor (Roche), and 1x PhosSTOP (Roche)). Lysates were cleared via 14000x g centrifugation at 4°C, and split evenly for RNA vs protein processing. RNA extraction, cDNA synthesis, and qRT-PCR was carried out as described above. Lysates for protein work were mixed 1:1 with 2x Laemmli sample buffer, and boiled for 1 min. Proteins were resolved via SDS-PAGE, blotted to Hybond ECL membrane (GE Healthcare), and blocked in 5% milk fat. PHB variants were detected via anti-GFP primary antibodies (Rockland Immunochemicals; 1:500 dilution) and anti-rabbit horseradish peroxidase-conjugated secondary antibodies (Jackson Immunoresearch; 1:5000 dilution). Detection of secondary antibodies was performed with SuperSignal West Pico Chemiluminescent Substrate (ThermoFisher Scientific). Blots were scanned and quantified using ImageJ. Four independent biological replicates were performed, each with two technical replicates.

### Protein immunoprecipitation and mass spectrometry

One gram of 24 hrs estradiol-induced seedlings was flash frozen, ground under liquid nitrogen, and extracted in 4 ml of freshly prepared Extraction Buffer (50 mM HEPES pH 8, 150 mM NaCl, 0.3% NP-40, 1x cOmplete ULTRA protease inhibitor - EDTA (Roche), 1mM PMSF, 50 uM bortezomib). Lysates were cleared via 14000x g centrifugation at 4°C, pre-cleared with Protein A Dynabeads, and incubated for 1 hr with 5μl anti-GFP antibody (abcam ab290). PHB variant complexes were captured using 25 μl Protein A Dynabeads and washed 4x with freshly prepared Wash Buffer (50mM HEPES pH 8, 450mM NaCl, 0.1% NP-40, 1x cOmplete ULTRA protease inhibitor - EDTA (Roche), 1mM PMSF, 50uM bortezomib).

After the final wash, beads were flash frozen in liquid nitrogen and trypsinized via standard protocols (Promega). Peptides were labeled with 8-plex iTRAQ^*66*^, and injected into the Orbitrap Velos Pro mass spectrometer. Protein identification and quantification was carried using Mascot 2.4^*67*^ against the UniProt *Arabidopsis* sequence database. Enrichments over the uninduced control were calculated for each replicate, mean enrichment was then calculated across the top five replicates, and a Student’s t-test applied to determine significance. Known false positives were eliminated from subsequent analyses^*68*^. Interacting partners for PHB-SDmut and PHB-Delta were identified using a similar workflow.

### Protein purification, circular dichroism, and membrane-overlay assays

Wild-type or START mutant variants (496-1137bp from the start codon) were cloned downstream of maltose binding protein (MBP), in a modified pET28b vector^*69*^, via Gibson cloning (ThermoFisher). Proteins were induced with 100 μM IPTG at 12°C for 16 hrs and purified on a Ni^2+^ column according to manufacturer protocols (Qiagen). Proteins were eluted using 125 mM imidazole, desalted and concentrated on a 10 MWCO column (Amersham) into NaCl-free IP buffer (1x PBS pH7.4, 5% glycerol), and quantified next to BSA on a 10% SDS gel.

Circular dichroism was performed with the Jasco J-815 Spectrometer (OSU Biophysical Interaction & Characterization Facility) using 1mg/ml of purified MBP or MBP-START variant proteins, then corrected and converted to molar ellipticity in Excel (Microsoft). Scans were limited to a range of 190-250nm to increase resolution and data were plotted in GraphPad (Prism).

Membrane-overlay assays were done according to manufacturer protocols (ThermoFisher Scientific; P23751). In brief, membranes were blocked for 1h at RT (PBS, 0.1% Tween20, 3%BSA fatty acid free). 150ug of wild-type START purified protein was added to the membrane in 5ml blocking buffer and incubated for 1h with gentle shaking. Membranes were washed three times with PBS-T and incubated with 1:2000 anti-His-tag antibody (Abiocode M0335-1) for 1h with gentle shaking at RT. Membranes were washed three times and incubated with 1:2000 anti-mouse 2° antibody (ab6789) for 1h. Bound proteins were detected using Hyper HRP Substrate (Takara). PtIns(3,5)P_2_ Grip protein (P-3516-3-EC, MoBiTec), detected via anti-GST antibody (Sigma Aldrich, A7340), served as a positive control (not shown).

#### Liposome production and dynamic light scattering

Total lipids from 35-40g of 21d old *Arabidopsis* seedling tissue were extracted using Bligh-Dyer^*70*^. Lipids were dried under argon and resuspended in 100% ethanol to 100mg/ml. Liposomes were generated by rapid injection of ethanol-suspended lipids into IP buffer (1x PBS pH7.4, 150mM NaCl, 5% glycerol). Liposome size range was determined by dynamic light scattering using the Zetasizer Nano (Malvern).

#### Lipid immunoprecipitations and mass spectrometry

To identify lipids bound by the START domain in bacteria, lipids from purified recombinant MBP and MBP-START proteins were immediately extracted using Bligh & Dyer^*70*^, dried under argon, and resuspended in chloroform. Lipids were identified using LC-MS performed at the Institut fur Molekulare Physiologie und Biotechnologie der Pflanze (IMBIO) at the University of Bonn, Germany. In brief, lipids were resuspended in 200ul Q-ToF solvent was added (methanol/chloroform/300 mM ammonium acetate, 665:300:35, v/v/v)^*71*^, and measured using nanoflow direct infusion Q-TOF MS/MS (as described in ^*72*^).

To identify lipids bound by the START domain in plant lipid mixtures, 1mg of recombinant MBP or MBP-START proteins were incubated with 6mg of liposomes in IP buffer (1x PBS pH 7.4, 150mM NaCl, 5% glycerol) overnight at 4 degrees. IPs were performed in quintuplicate. Proteins were repurified using nickel affinity chromatography, extensively washed, and eluted using 125mM imidazole. Lipids were extracted using Bligh & Dyer^*70*^, dried under argon, and resuspended in chloroform. Lipids were identified using LC-MS performed at the Kansas Lipidomics Research Center Analytical Laboratory (KLRC; Kansas State University) and the Nutrient & Phytochemical Analytics Shared Resource (NPASR; Ohio State Comprehensive Cancer Center). KLRC used a Waters Xevo TQS mass spectrometer adjusted for SPLASH response factors. 400ul volumes were injected, ES+ ionization mode was used for all detection of all compounds except lysoPG which used ES-, and data presented are nmol per mg dry weight and normalized to mols of protein per IP. NPASR data was using a SelexION/QTrap 5500 (Sciex) Lipidyzer for CER, LPC, LPE, PC, PE, and SM and a 6550 QTOF MS (Agilent) for FFAs. 150ul volumes were injected, ESI-ionization mode was used, and data presented are normalized to mols of protein per IP.

We note that KLRC instrument acquisition and lipidomics method development was supported by National Science Foundation (EPS 0236913, MCB 1413036, MCB 0920663, DBI 0521587, DBI1228622), Kansas Technology Enterprise Corporation, K-IDeA Networks of Biomedical Research Excellence (INBRE) of National Institute of Health (P20GM103418), and Kansas State University. We also note that research reported in this publication was supported by The Ohio State University Comprehensive Cancer Center and the National Institutes of Health under grant number P30 CA016058.

## Data availability

All data is available in the main text or the Supplemental Data section.

## Acknowledgements

We are grateful to D. Skopelitis, L. Joshua-Tor, D. Pappin, and K. Rivera from Cold Spring Harbor Laboratory, and Diana Vranjkovic and Phillipp Johnen from the University of Tuebingen for technical support. We are grateful to Dr. Zachary Schultz for assistance with dynamic light scattering assays, and Dr. Alicia Friedman for assistant with circular dichroism. We thank Dr. Ruth Welti and Mary Roth at the KLRC, and Dr. Ken Riedl at the OSUCCC NPASR, for lipidomic analyses. We thank Dr. Mark Stahl at the Center for Plant Molecular Biology at the University of Tübingen for assistance with LC-MS. We also thank Dr. Doris Wagner for critical comments on the manuscript. Support for this work came from National Science Foundation grants IOS-1022102 and IOS-1355018, as well as the Alexander von Humboldt Professorship, to M.T. Support for this work also came from National Science Foundation grant IOS-2039489 to A.Y.H.

## Corresponding authors

Correspondence to Aman Y. Husbands or Marja C. P. Timmermans

## Contributions

A.Y.H and M.T conceived of the research direction and experiments; A.Y.H. performed experiments except: protein purification by A.F., LC-MS by KLRC and NPASR centers, additional ChIP and protein work by C.E.D. and A.S.H, and SiMPull by V.A. under the guidance of T.H. A.Y.H and M.T. wrote the manuscript.

## Supplementary Information

**Supplementary Fig. 1.**
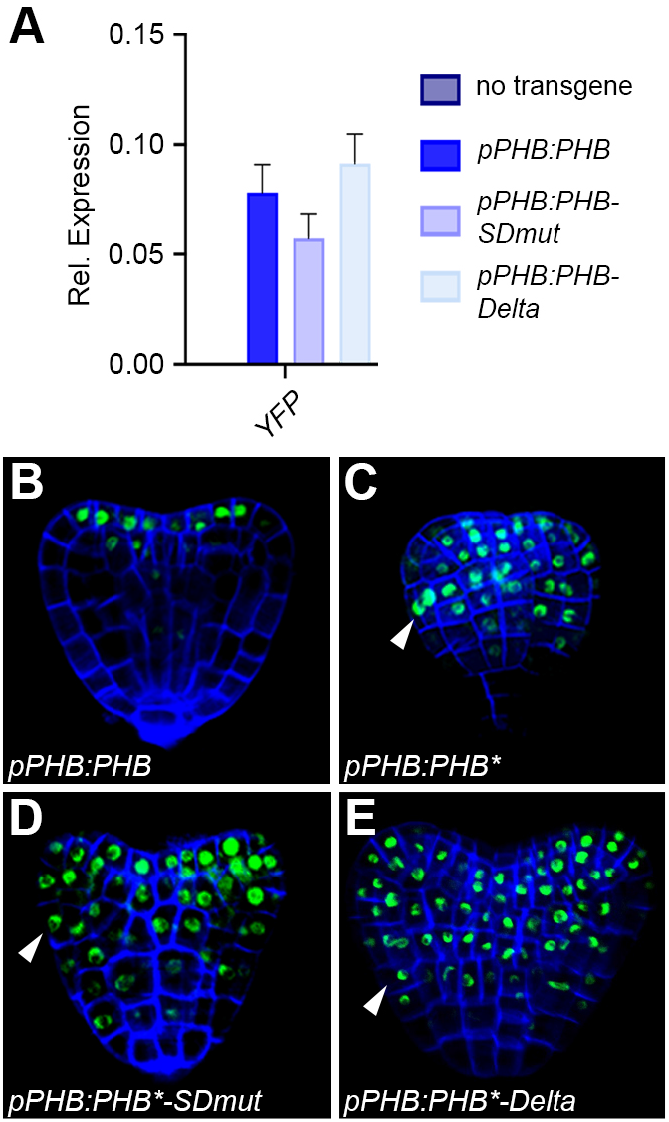
The disparate behaviors of *PHB* variants are not explained by differences in transcript accumulation, nuclear protein localization, or ectopic expression in the absence of miR166-regulation. **A**, *PHB-YFP, PHB-SDmut-YFP*, and *PHB-Delta-YFP* transcripts accumulate to equivalent levels in *phb phv cna* complementation lines, although only the *pPHB:PHB-YFP* transgene complements the triple mutant phenotype (**Fig. 1B**). **B**, miR166 activity limits PHB expression to the adaxial side of developing cotyledons and to incipient vasculature in heart-stage embryos. **C-E**, Loss of miR166-regulation leads to ectopic accumulation of PHB, PHB-SDmut, and PHB-Delta throughout the embryo (white arrowheads). All three PHB variants localize to the nucleus. Development is delayed by ectopic accumulation of PHB (**C**), but not by PHB-SDmut or PHB-Delta (**D, E**). Cell walls are marked by fluorescent brightener 28. A minimum of 20 individual transgenic lines were screened and representative images shown here. Unlike START-deleted HD-ZIPIV proteins^*32,33*^, all PHB variants are easily detectable and accumulate to qualitatively similar levels in all lines.

**Supplementary Fig. 2.**
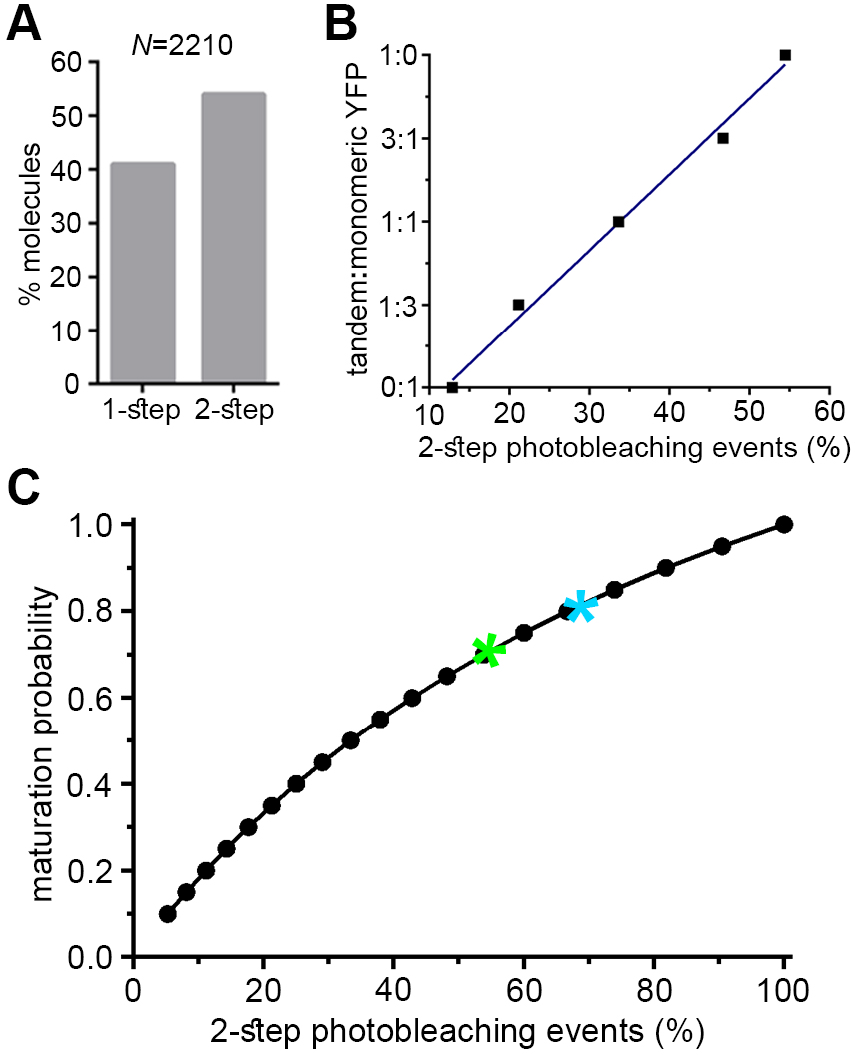
Citrine YFP maturation probability in *Arabidopsis* and construction of a SiMPull calibration curve. Fluorophore photobleaching is marked by an abrupt decrease in fluorescence intensity, allowing the number of dimers in a population to be inferred by the frequency of two-step photobleaching events, provided the maturation probability of the fluorophore in the system is known. **A**, SiMPull and photobleaching analyses show 41% one-step vs 54% two-step photobleaching events with *Arabidopsis* lysates containing dimers of citrine YFP. **B**, *Arabidopsis* lysates containing dimeric or monomeric citrine YFP were mixed at dimeric:monomeric ratios of 0:1, 1:3, 1:1, 3:1, or 1:0. Using SiMPull and photobleaching analyses, a calibration curve (y = 2.28x - 27; R^2^ = 0.992) was constructed to relate two-step photobleaching events to frequency of dimers in the population. **C**, Binomial probability modeling predicts a maturation probability of 0.7 for citrine YFP in *Arabidopsis* (green asterisk); maturation probability of citrine YFP in mammalian systems (cyan asterisk) is plotted for comparison^*44–46*^.

**Supplementary Fig. 3.**
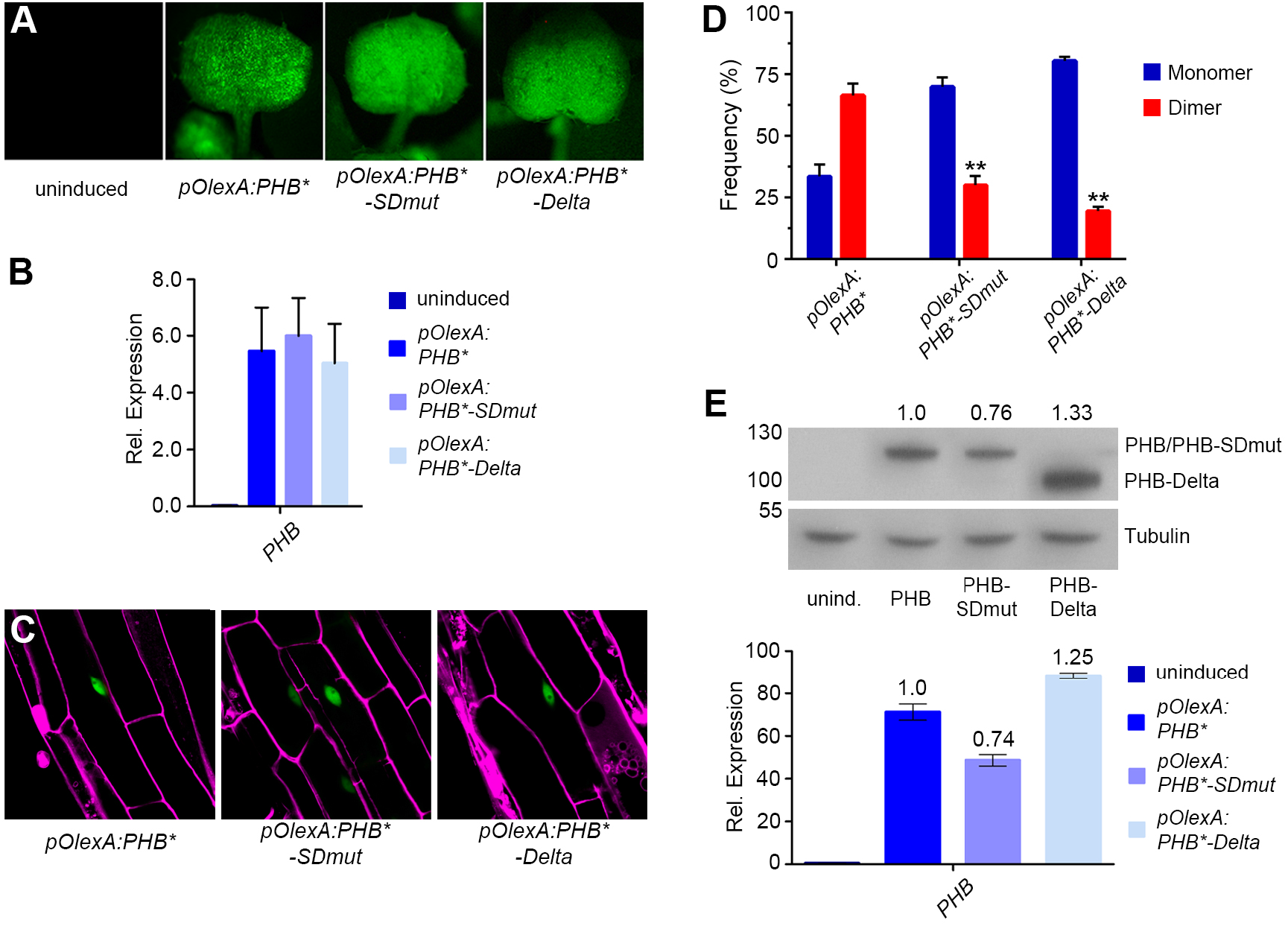
Subcellular localization, dimerization frequency, and protein stability are unaffected upon estradiol-induced overexpression of PHB variants. **A, B**, 24 hrs post-estradiol-induction, *pOlexA:PHB*-YFP*, *pOlexA:PHB*-SDmut-YFP*, and *pOlexA:PHB*-Delta-YFP* seedlings show near equivalent, strongly-elevated expression of each respective PHB variant, as measured by fluorescent imaging (**A**) and qRT-PCR (**B**). **C**, PHB variants produced in estradiol-inducible lines retain their nuclear-localization. Cell walls are marked by propidium iodide (magenta). **D**, SiMPull with extracts from 24 hrs estradiol-induced *pOlexA:PHB*-YFP*, *pOlexA:PHB*-SDmut-YFP*, and *pOlexA:PHB*-Delta-YFP* seedlings shows dimerization frequencies of ~67% for PHB, ~31% for PHB-SDmut, and ~19% for PHB-Delta, comparable to the frequencies observed for lines expressing these variants from the native *PHB* promoter (**Figs. 2B** and **5A**). **E**, Levels of PHB variant protein and RNA – extracted simultaneously from 24 hrs estradiol-induced seedlings – quantified using Western blotting (top) and qRT-PCR normalized to *PHB* levels (bottom), respectively, shows RNA levels correlate with protein levels for each PHB variant, indicating START mutation or deletion does not dramatically affect protein stability. Data was generated for four independent replicates. In the interest of clarity, one biological replicate, comprised of two technical replicates, is shown. \*\**P* ≤ 0.01, Student’s *t*-test.

**Supplementary Fig. 4.**
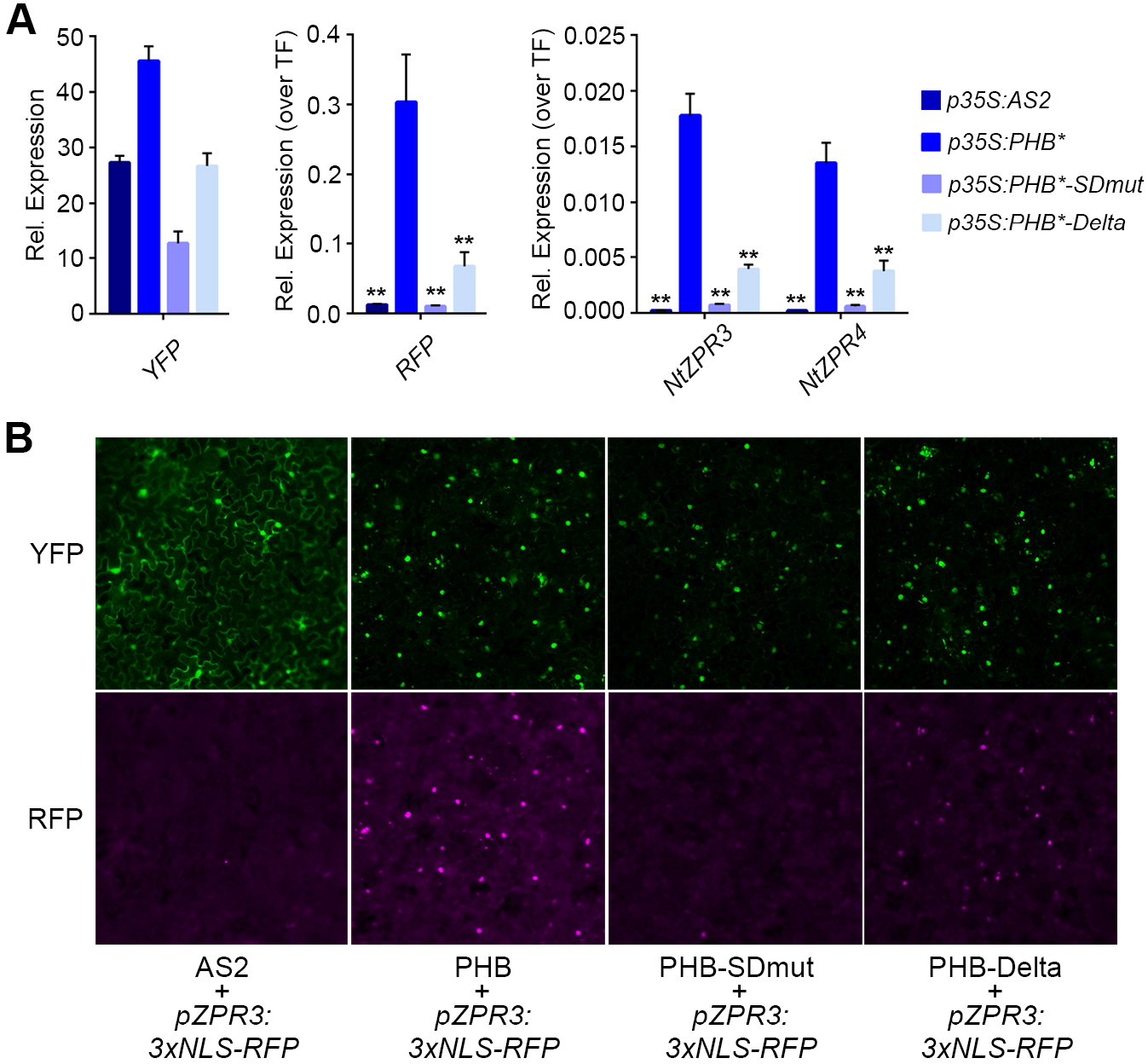
The START domain increases transcriptional potency. **A**, *NtZPR3, NtZPR4*, and *RFP* levels in tobacco leaves 24 hrs after transfection with *p35S:AS2-YFP, p35S:PHB*-YFP, p35S:PHB*-SDmut-YFP*, or *p35S:PHB*-Delta-YFP*, and a *pZPR3:3x-NLS-RFP* reporter. PHB-Delta is ~3-fold weaker at activating transcription than PHB, while PHB-SDmut and the AS2 negative control fail to activate these HD-ZIPIII targets (*n* = 3 biological replicates). Molecular behaviors of PHB variants in tobacco thus mirror that of *Arabidopsis* (**Figs. 3** and **5C-D**). **B**, Consistent with quantitation in **A**, fluorescent imaging shows *pZPR3:3x-NLS-RFP* reporter activity is strongest in leaves accumulating PHB, weaker in leaves accumulating PHB-Delta, and not detectable in leaves accumulating PHB-SDmut or AS2. \*\**P* ≤ 0.01, Student’s *t*-test.

**Supplementary Fig. 5.**
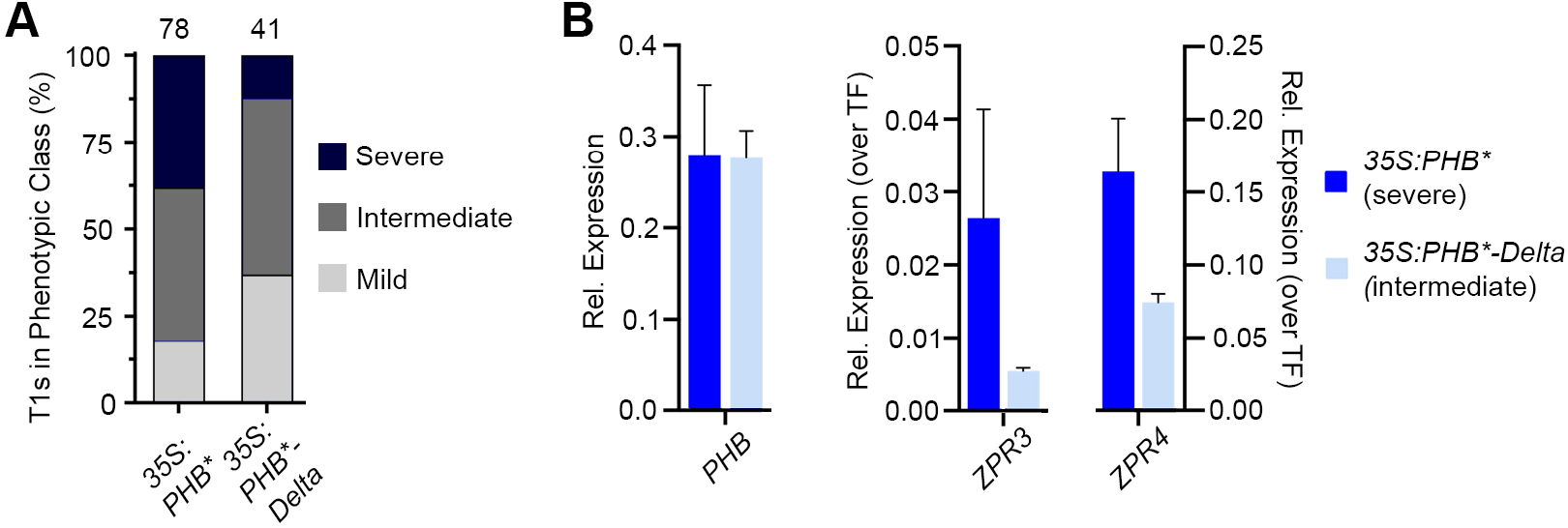
PHB-Delta dimers have reduced transcriptional potency. **A**, *p35S:PHB*-Delta* primary transformants show intermediate and severe gain-of-function phenotypes, consistent with this PHB variant retaining partial function. Overall, the phenotypes of *p35S:PHB *-Delta* transformants are less severe than those observed for *p35S:PHB\** transformants (*n* above bar). Note that, as the *35S* promoter is less active early in development, seedling phenotypes are generally weaker than those seen following transformation of *pPHB:PHB\** (**Fig. 1C**). **B**, Compared to lines with comparable overexpression of *PHB* (left), PHB-Delta lines exhibit weaker phenotypes and accumulate lower levels of their *ZPR3* and *ZPR4* targets (right).

**Supplementary Fig. 6.**
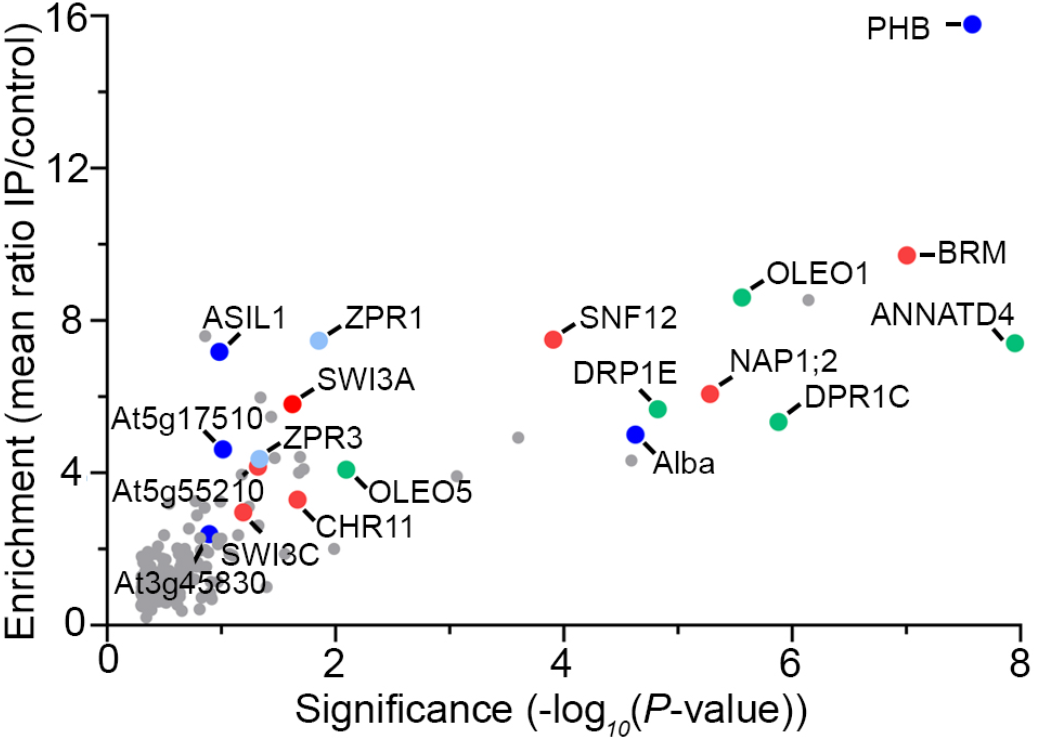
PHB binds multiple classes of interacting partners in a START-independent manner. One-sided volcano plot showing enrichment values and corresponding significance levels for proteins co-immunoprecipitating with PHB (*n* = 5 biological replicates). These include multiple components of the SWI/SNF chromatin-remodeling complex (red), as well as ZPR proteins known to interact with HD-ZIPIII proteins (lavender). Other enriched factors include transcriptional regulators (blue) and lipid-interactors (green). PHB-SDmut and PHB-Delta share this spectrum of interaction partners. See also **Tables S1** and **S2**.

**Supplementary Fig. 7.**
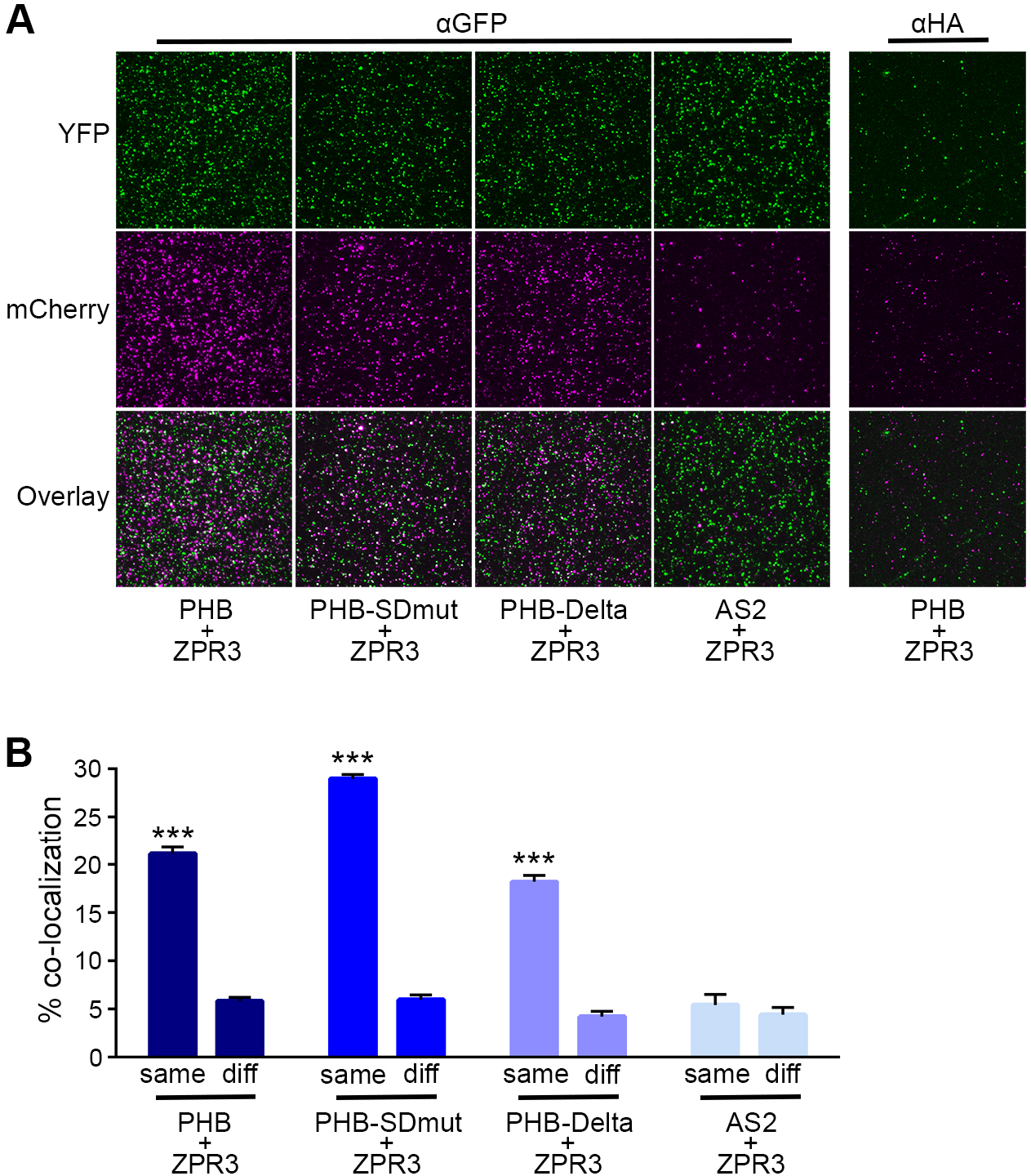
PHB - ZPR3 interaction is not affected by START mutation or deletion. **A**, Representative TIRF images from SiMPull using tobacco leaf lysates co-expressing PHB-YFP, PHB-SDmut-YFP, or PHB-Delta-YFP with ZPR3-mCherry, or from lysates co-expressing AS2-YFP with ZPR3-mCherry. Overlaid TIRF images of YFP (green) and mCherry (magenta) channels show pronounced colocalization of all PHB variants with ZPR3 (white spots in Overlay panels); AS2 and ZPR3 colocalization, however, is indistinguishable from α-HA background, indicating these proteins do not interact. More details on SiMPull are in ^*18,44–46*^. **B**, Co-localization frequencies for each PHB variant with ZPR3 are significantly higher for overlaid TIRF images taken from the same region of SiMPull slides (same) than from random, non-overlapping regions (diff), or from the AS2 and ZPR3 negative control. Colocalization frequencies (means ± SE) are calculated from at least 30 images, from three independent biological replicates. Student’s t test: ***P < 0.001.

**Supplementary Fig. 8.**
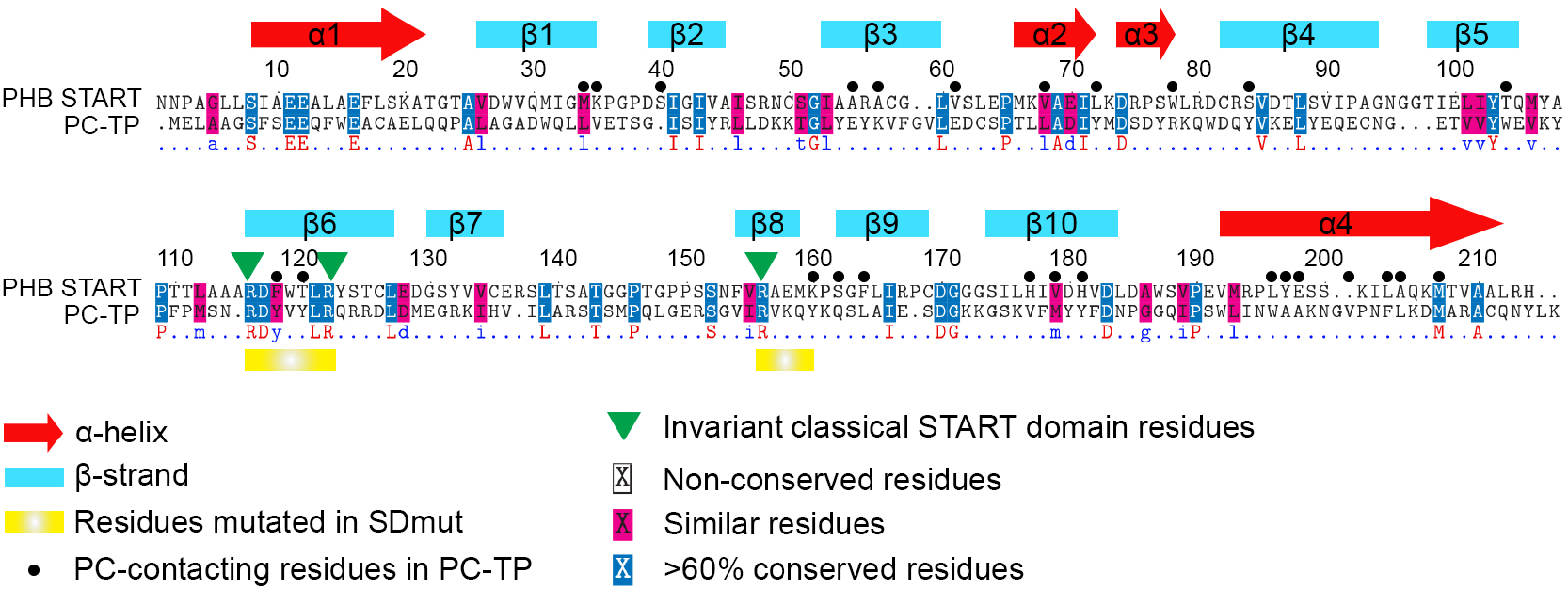
The PHB START domain most closely resembles PC-TP. Pairwise sequence alignment of the PHB START domain and PC-TP with positions of key features including α-helices (red arrow), β-sheets (blue boxes), PC-contacting residues (black dots), invariantly-conserved arginine residues (green arrowheads), and residues mutated in PHB-SDmut (yellow boxes) indicated. Two regions surrounding three invariantly conserved arginine residues^*8*^ were selected for mutation; specifically, RDFWTLR was changed to GAVVGAG and RAEMK to VAAGV.

**Supplementary Fig. 9.**
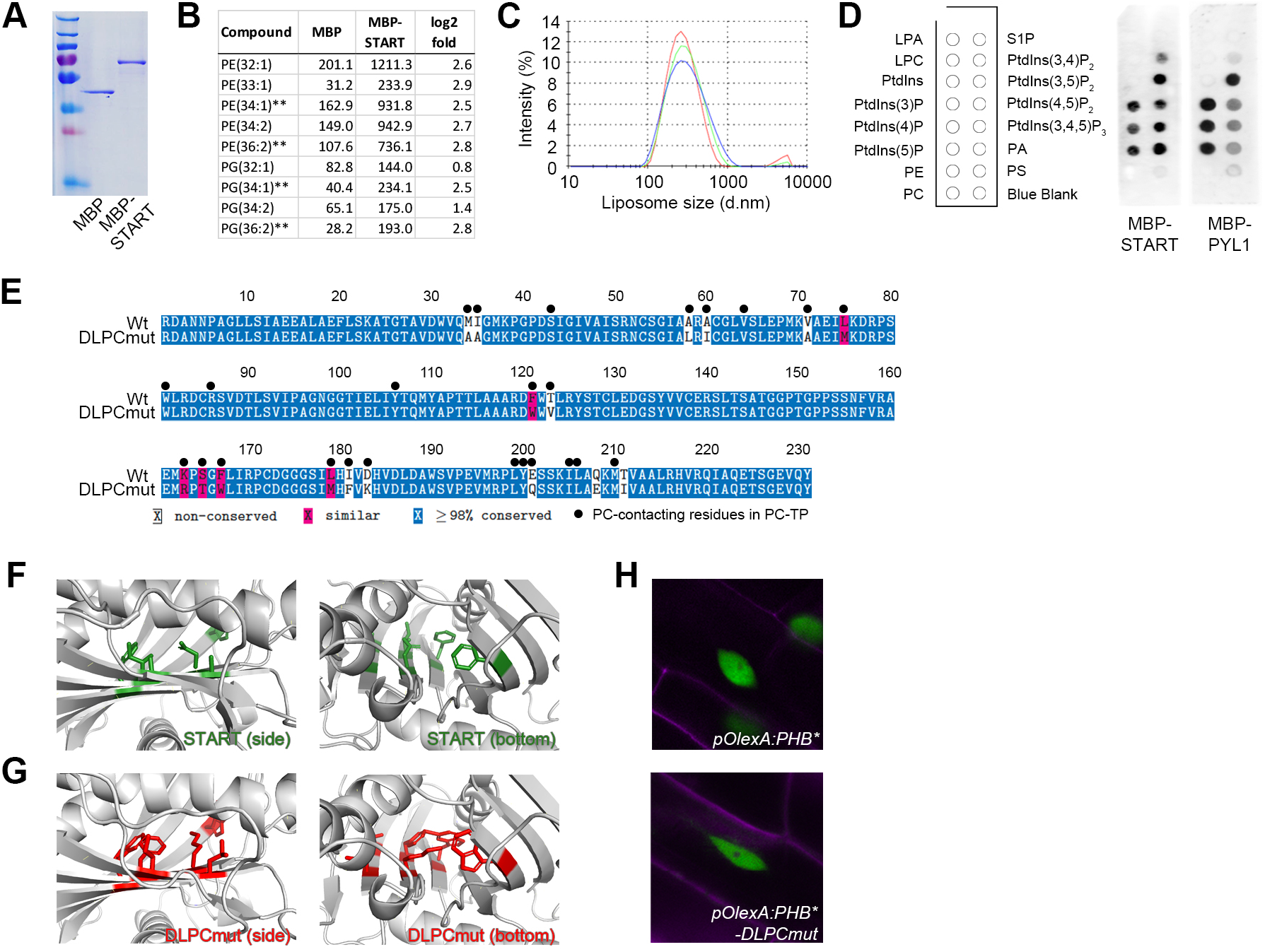
The START domain binds phospholipids, and mutations in putative ligandcontacting residues do not affect folding or nuclear localization. **A**, Purified MBP and MBP-START recombinant proteins. **B**, LC-MS analysis of these recombinant proteins shows enrichment of multiple PE and PG species in MBP-START. Raw signal intensities are reported, ** indicates species preferentially bound by PC-TP^*31*^. **C**, Differential Light Scattering showing liposomes made from *Arabidopsis* total lipid extracts are uniformly sized (~300nm diameter). **D**, Membrane-overlay assays show binding to PE and PC does not occur at the surface of the START domain. Similar assays using the PYL1 StARkin domain indicate the surface of both domains interacts with several negatively charged lipids. **E**, Alignment showing residues mutated in the DLPCmut START variant. Residues were chosen based on information from PHB START homology modeling and known PC-containing residues of PC-TP (**Fig. S8**). All substitutions were chosen to minimize effects on secondary structure. **F, G**, Homology models of wildtype and DLPCmut START domains (structures in grey; side view left panels; bottom view right panels). Unlike wild-type residues (side chains in green), six DLPCmut substitutions (A58L, A60I, F121W, F167W, I181F, D183K; side chains in red) partially occlude the binding pocket to compete with entry of ligands. **H**, PHB-DLPCmut protein retains a nuclear-localization. Cell walls are marked by propidium iodide (magenta).

